# Large-scale automated detection reveals pervasive sex imbalance in biomedical research

**DOI:** 10.64898/2026.07.13.738332

**Authors:** Lydia E. Valtadoros, Parker Hicks, Hao Yuan, Mansooreh Ahmadian, Kayla A. Johnson, Arjun Krishnan

**Affiliations:** Department of Biomedical Informatics, University of Colorado Anschutz Medical Campus, Aurora, CO; Genetics and Genome Sciences Program, Michigan State University, East Lansing, MI; Ecology, Evolution, and Behavior Program, Michigan State University, East Lansing, MI; Department of Biostatistics and Informatics, University of Colorado Anschutz Medical Campus, Aurora, CO

**Keywords:** sex as a biological variable, natural language processing, data reuse

## Abstract

Sex is a critical biological variable that impacts disease risk, progression, and treatment response across virtually every organ system. However, decades of biomedical research have relied primarily on male study subjects, leaving large gaps in our understanding of female-specific disease biology. Quantifying the extent of this imbalance across thousands of disease areas and millions of publicly available biological samples has remained computationally intractable. Here, we present a multimodal computational framework that infers the biological sex of ∼230,000 publicly available human transcriptome samples and links inferred sex labels to disease terms extracted from ∼9,000 associated study records and ∼5,000 publication abstracts to quantify sex imbalance at scale. Applying this approach revealed that the majority of disease terms with the largest research-derived sex imbalance are skewed toward male representation, including areas with no known biological justification for that imbalance. After adjusting for global sex-specific disease prevalence to isolate biologically unjustified imbalance, up to 58% of all disease terms showed male-leaning association. Diseases including glioblastoma, cirrhosis, idiopathic pulmonary fibrosis, and schizophrenia emerged as critically understudied in females despite affecting both sexes comparably. These findings provide a principled, data-driven basis for prioritizing compensatory research efforts and offer a reusable framework for ongoing monitoring of sex representation in the biomedical literature.

**Highlights:**

- Skewed male and female study subject representation in biomedical research is the result of decades of studies conducted without adequate female representation.
- We developed an automated, multimodal framework to estimate the sex imbalance across thousands of disease terms using metadata from ∼230,000 transcriptomics samples and their associated ∼9,000 studies and ∼5,000 publications.
- Our approach identifies non-sex-specific disease research areas that have been studied using an unbalanced sex demographic. These areas need compensatory and balanced studies to understand sex differences.

## Introduction

Female study subjects in human, animal, and cell-culture studies have been inadequately included in decades of research in multiple biomedical fields [1,2]. The historical underrepresentation of female study subjects from biomedical research has introduced an imbalance in male- and female-specific knowledge in different areas where certain diseases are detrimentally understudied in one biological sex. While some diseases are sex-specific (e.g., prostate cancer only affecting males), others have no known biological justification to have significant sex representation imbalance, as they affect both females and males (e.g., cardiovascular disease, lung cancer). It is vital to identify these areas and quantify their sex imbalances for guiding compensatory research efforts, as a lack of rigorous research using female models has serious public health consequences [3–8].

Existing efforts have previously quantified sex imbalances across specific biomedical research areas, yet these efforts are on a smaller scale and are usually done manually [6,9–13]. Further, the rapid pace at which scientific work is being conducted makes it impossible to investigate more than a few journals, years, or articles by hand. The exponential increase of primary articles and the biological samples that are collected in the associated studies [14] introduces a need for automated, large-scale identification of research areas that have sex imbalances. To the knowledge of the present study, few studies have quantified sex imbalance on a large-scale level.

Existing methods share three limitations that constrain their reach. First, approaches relying on explicitly reported sex counts [15] are limited by incomplete and inconsistent reporting. Approximately half of clinical trials posted at ClinicalTrials.gov have no corresponding publication [16], and demographic data often differs between trial records and papers [16]. Second, approaches drawing on repository-provided sex annotations [17] are constrained by the fact that sex labels are absent for the majority of publicly available samples [18]. Third, even approaches that predict sex labels directly from sample expression data [19] have used exact text matching to extract disease terms from metadata, missing synonyms, abbreviations, and related terminology that named-entity recognition methods can capture. Taken together, prior work has assessed sex imbalance either on a small scale, or on a large scale but in a limited disease space, or without leveraging the disease-rich metadata embedded in publication abstracts and study descriptions associated with publicly available transcriptomic samples.

Here, we present a large-scale and data-driven analysis of sex representation imbalance in disease studies by integrating publicly available transcriptomes and their associated metadata together. We avoid annotation incompleteness by inferring sex (for ∼230,000 human transcriptomic samples) directly from expression profiles rather than relying on metadata annotations. We capture disease information across two complementary metadata modalities by combining sex-labeled transcriptomes with NLP-derived disease terms from both study metadata (∼9,000 studies) and publication abstracts (∼5,000 publications). And, we separate research imbalance from biologically justified imbalance, which represents a genuine gap in female representation, by adjusting inferred imbalance values for sex-specific global disease prevalence. Using this multi-modal approach, we identify groups of diseases and individual disease terms with existing research skewed toward one sex that require further, compensatory attention to improve our understanding of sex-driven effects in various biomedical research areas.

## Methods

The goal of our work was to identify disease research areas understudied in females or males, and quantify that imbalance, using thousands of publicly available human transcriptomic samples and their associated publication and study metadata. We first inferred the sex of each sample in our transcriptome compendium and extracted relevant disease terms from the text metadata associated with our samples. We combined this information to identify individual disease terms and groups of terms that have been studied using an unbalanced sex demographic, even after accounting for sex-specific global disease prevalence (**Figure 1**).

**Figure 1.**
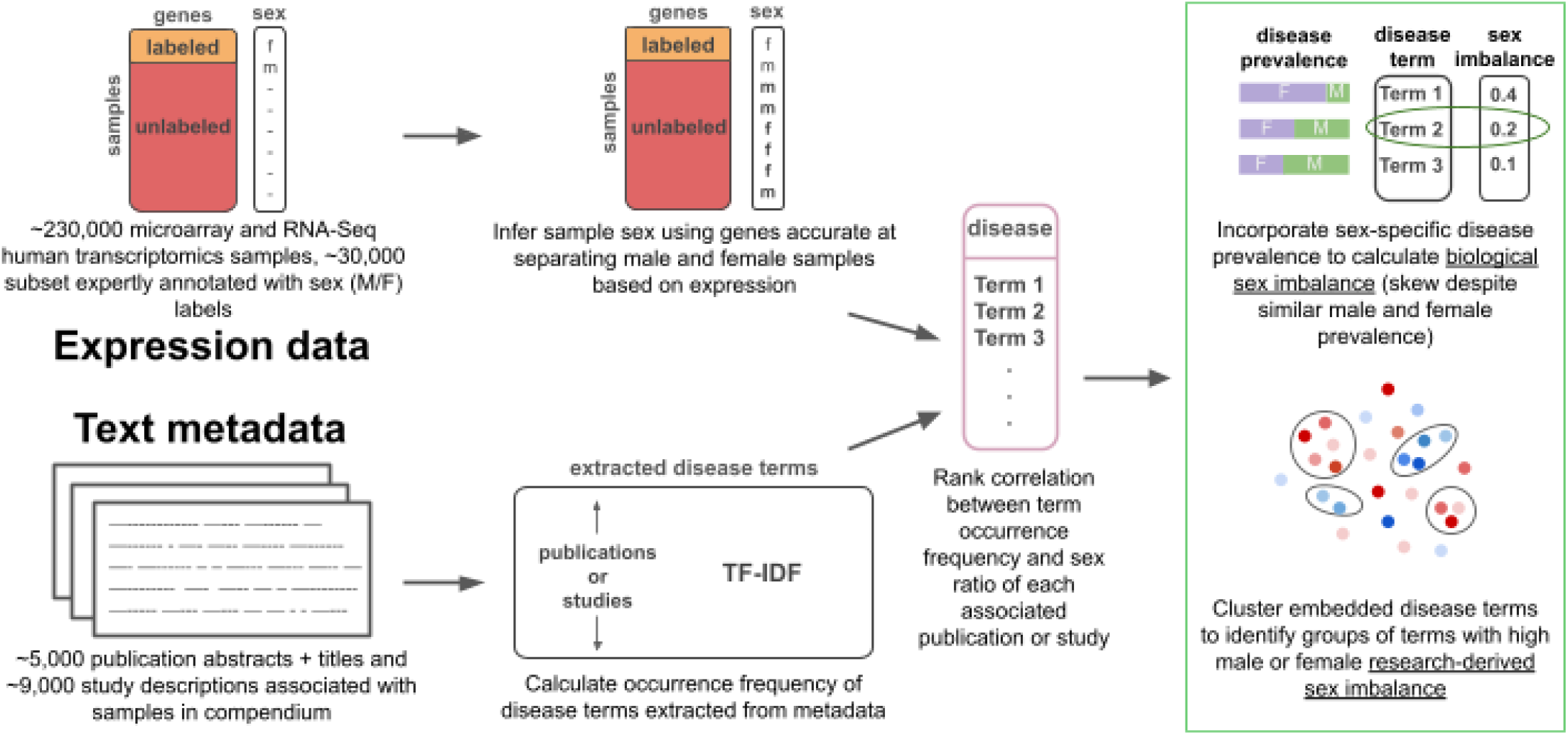
Expression data for ∼230,000 human transcriptomics samples was used to infer sex labels for all samples in the compendium. Sex label results were combined with the occurrence frequency of biomedical disease terms extracted from publication and study metadata associated with the transcriptomics samples. The correlation between the occurrence frequency of each term with the sex ratio of female to male samples in each publication or study represents the research-derived sex imbalance of the disease terms, which was adjusted with sex-specific global prevalence to quantify biological sex imbalance.

### Data collection

We downloaded the expression levels and sample descriptions for 229,528 human transcriptomic samples from the Gene Expression Omnibus [20] (GEO) and refine.bio [21]. Microarray samples were downloaded from GEO, which included all human samples measured on the *Affymetrix Human Genome U133 Plus 2*.*0 Array* platform. RNA-Seq samples included all human samples available in refine.bio at time of data accession.

Expression data for microarray samples was downloaded from GEO as raw CEL files, which were processed with background subtraction, quantile transformation, and summarization using fRMA based on custom CDF mapping probes to Entrez gene IDs [22]. RNA-Seq sample expression data was downloaded as Salmon output files. Samples with over 50% of zero counts were removed. RNA-Seq expression values were quantified in transcripts per million (TPM). Only genes measured for both technologies were included, resulting in 18,478 genes [22]. There were 124,748 microarray samples and 104,780 RNA-Seq samples, totaling 229,528 human transcriptomic samples used in analysis.

### Gold-standard sex and age labels

To create our gold-standard dataset for inferring sample sex, we extracted sex and age labels for 30,339 samples (16,107 microarray, 13,733 RNA-Seq) from sample descriptions through text-mining and manual curation [22,23]. Age labels were assigned by categorizing samples into eleven previously defined groups based on hormone signatures across the human lifespan [22], and were leveraged to identify genes indicative of sample sex for samples of all age groups. Genes with high accuracy in distinguishing male and female samples across all age groups were used for inferring the sex labels of all samples.

We downloaded the sample descriptions for microarray data from GEO and verified the text-matched sex and age labels by reviewing the sample descriptions. For RNA-Seq data, we downloaded sample-level sex and age annotations from metaSRA [24] version 1.8. We then used ffq [25] to retrieve sample and study descriptions from Sequence Read Archive (SRA) [26]. We mapped the corresponding sex and age labels to each run identifier in refine.bio and verified the labels by comparing the annotations to the descriptions.

### Inferring sample sex labels

We inferred the sex for all of our samples to quantify the number of male and female samples associated with the studies in our corpus. All sex inference analyses were performed separately for microarray and RNA-Seq samples. First, we used our gold-standard subset annotated with sex and age labels to find genes best at separating male and female samples based on an expression threshold specific to each gene. We recorded this threshold and the balanced accuracy for each gene (i.e., how well a gene separated male and female samples), as well as which sex was associated with higher expression. We performed this process for each of the eleven predetermined age groups (**Figure 2a**). We selected the nine genes with the highest average balanced accuracy across all age groups for microarray and RNA-Seq samples separately (**Figure 2b**). All genes used had a balanced accuracy above 0.82 in all age groups. Including additional genes beyond the top nine decreased the average balanced accuracy below 0.75, so only the top nine were used.

**Figure 2.**
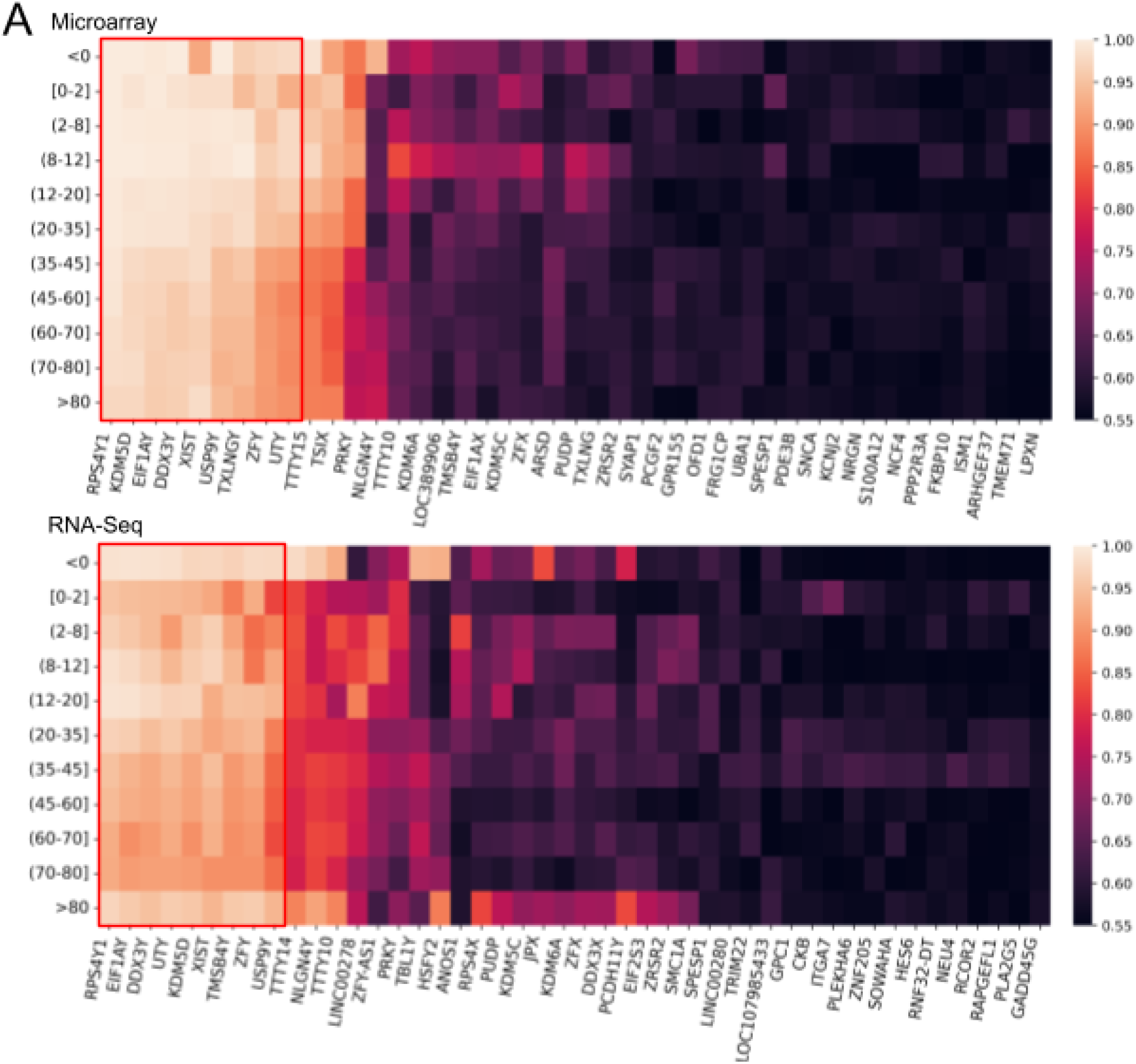
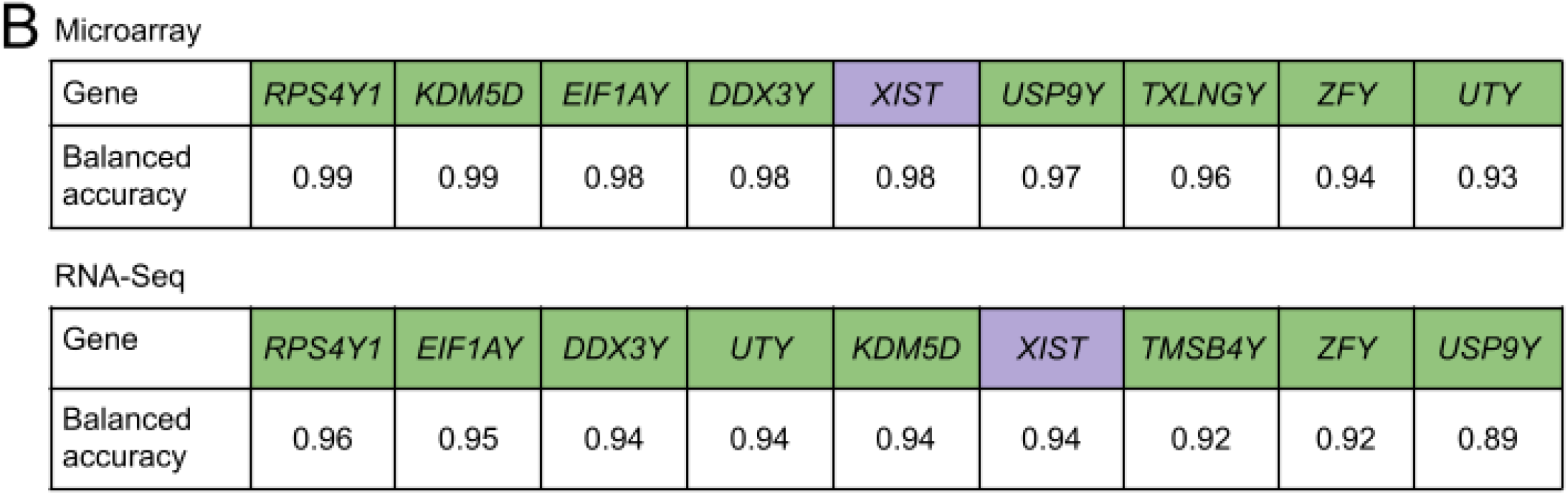
Heat map of gene balanced accuracies in all eleven age groups for genes with highest accuracy averaged across all age groups. Genes in the red box were used for predicting microarray (top) and RNA-Seq (bottom) samples (A). Balanced accuracy of genes used for predicting the sex of microarray (top) and RNA-Seq (bottom) samples (B). Genes highlighted green are located on the Y chromosome, genes highlighted purple are located on the X chromosome.

The majority of genes identified as indicative of sex in both microarray and RNA-Seq analyses are located on the Y chromosome (*RPS4Y1, KDM5D, EIF1AY, DDX3Y, USP9Y, UTY, ZFY* for both technologies, *TXLNGY* only for microarray and *TMSB4Y* only for RNA-Seq). Only one gene indicative of sex for both technologies is located on the X chromosome (*XIST*).

We inferred the sex of of all ∼230,000 samples in our compendium by comparing a sample’s expression level for each predicting gene to the expression threshold for that gene, as the expression of genes located on sex chromosomes clearly indicate female or male classification [27,28]. For example, if the sample expression for gene 1 was above gene 1’s threshold, and expression levels higher than the threshold were associated with male samples, gene 1 indicated the sample as male. This process was completed for all nine genes, so the inferred sex for each sample was the majority sex as indicated by the predicting genes.

To investigate the reliability of our inferred labels, we assessed the quality of our samples by comparing the expression of each gene for a given sample to the average expression for all samples of the same inferred sex. First, we recorded the 25th percentile expression for each gene (first subsetting for genes expressed at similar levels in female and male samples) across all samples of the same sex. Then, for each sample, we compared the expression of each gene to the 25th percentile expression level for that gene. We recorded the number of genes with low expression for each sample, such that samples with a high percent of low quality genes were deemed “poor quality.”

### Disease term extraction from publication and study metadata

We downloaded study descriptions to obtain study-level disease terms. There were 3,361 study descriptions associated with microarray samples and 5,347 study descriptions associated with RNA-Seq samples. For microarray samples, study descriptions were downloaded from SOFT files associated with each study in GEO. Study descriptions associated with RNA-Seq samples were accessed from SRA using ffq [25]. Disease terms were extracted from the descriptions using spaCy 3.7.4 [29] in conjunction with the “en_core_sci_md” model, which is designed to process biomedical text. There were 6,176 disease terms extracted from study metadata associated with microarray samples and 5,949 disease terms from RNA-Seq associated study metadata.

Publication metadata and disease annotations were accessed using PubTator3 [30], a text mining tool that provides pre-annotated biomedical entities for publication abstracts and titles. We first filtered available publications in PubTator3 for publications associated with samples in our compendium. For publications associated with microarray samples from GEO, PubMed was queried for each GEO study accession code using NCBI Entrez utilities [31] and all returned PubMed identifiers were linked to the corresponding GEO study. For publications associated with RNA-Seq samples from SRA, SRA records were first queried using NCBI Entrez utilities and parsed to identify corresponding GEO sample accession codes when available. These GEO sample accessions were subsequently linked to their corresponding GEO study accessions and publication identifiers. More than 25,000 articles were obtained from PubTator3 originally, but were reduced to 5,021 after filtering. Disease terms provided by Pubtator3 were used directly. Pubtator3 was not used for study descriptions because study metadata lacks the structured format that Pubtator3 requires. There were 2,971 terms from publications with microarray samples and 1,823 terms from publications with RNA-Seq samples.

### Quantifying research-derived sex imbalance

We first estimated “research-derived sex imbalance” based solely on the representation of female and male samples included in studies and publications without accounting for the disease prevalence in each sex. To do this, we first calculated a log-scaled term frequency-inverse document frequency (TF-IDF) score for each disease term (*t*) and document (*d*, study or publication metadata). A TF-IDF score quantifies the importance of a disease term within a document while reducing the influence of terms that occur frequently throughout the corpus. The TF-IDF of disease terms in each type of text was calculated as follows:

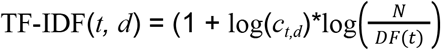

Where *c*_*t,d*_ represents the number of occurrences of term (*t*) in document (*d*), *DF(t)* represents the number of documents containing term (*t*) and (*N*) represents the total number of documents in the corpus. The corpus was defined separately for study descriptions and publication abstracts and titles. Higher TF-IDF values indicate disease terms that are both frequent within a specific document and relatively uncommon across the corpus.

Next, we calculated a “sex ratio” value for each publication by dividing the number of female samples by the total number of samples for each publication. We then calculated the Spearman rank correlation of the TF-IDF scores for each disease term with the publication sex ratio values. The correlation value for each term represents the research-derived sex imbalance of each term across the publication or study corpus. A high correlation magnitude indicated terms that are more frequent in metadata associated with only one sex; positive values indicated skew toward female overrepresentation and negative values indicated skew toward male. The same procedure was completed for each study and the disease terms extracted from study descriptions.

### Mapping extracted disease terms to Mondo identifiers

To enable comparisons of sex imbalance across related diseases, we first mapped disease terms to the Mondo Disease Ontology [32]. Specifically, a custom retrieval augmented generation (RAG) and OpenAI pipeline was developed to create this mapping. First, embeddings of the Mondo ontology terms was created using the sentence-transformers/nli-mpnet-base-v2 model [33]. Then, for each extracted disease term, a subset of possible high accuracy Mondo matches were identified by a RAG function. This function takes a query (“most related to the term: {disease term}”) and outputs the Mondo terms that best match the query based on the embeddings. From this subset, the OpenAI API was used to find the most accurate mapping from each original disease term to the possible Mondo terms. The instruction to the OpenAI API call asked the model to select the best match for the disease term based on the term name and knowledge of the model:

> “Here is a disease term: {disease_term}. Using the term name and your knowledge of the term, find the most similar disease out of the options provided in this string: {mondo_input}, each option is separated by ‘\n’. Prioritize exact term matches. If there is no match for the disease of the options, output ‘None’.
>
> Provide results in the format of one string with values separated by tab characters (\t):
>
> “{disease_term}\t’output’”
>
> With output being the most similar term from the options or ‘None’. Do not include any analysis of why the most similar term was chosen, just give the output in the format exactly as described. Make sure the most similar term you provide is actually from {mondo_input}, unless there is no match and the output is ‘None’”

Manual adjustments such as removing quotation marks, capitalizing terms, correcting spelling mistakes, and adding back missing parts of a truncated Mondo term were required for 20-40 terms (of ∼2,000-3,000 terms for study metadata and ∼6,000 terms for publication metadata, depending on technology). Upon completion of the automatic mapping and manual corrections, the map from the original disease terms extracted from the unstandardized, free-text metadata to Mondo ontology terms was complete.

### Clustering of Mondo-mapped disease terms

To identify groups of disease terms and assess the sex imbalance of disease areas, we first embedded the mapped Mondo disease terms. The Mondo Disease Ontology release version 2023-03-01 OBO file was converted into an undirected edge list using “is_a” and “part_of” relationships prior to embedding using the PecanPy [34] implementation of node2vec [35]. We constructed a k-nearest neighbors (KNN; k=3) graph from the node2vec embeddings and then clustered the disease terms using the Louvain community detection algorithm (*louvain_communities* function in NetworkX [36]). The same process was performed to cluster terms from the GEO-associated publications corpus, GEO study metadata, SRA-associated publications, and SRA study metadata. Depending on the metadata and disease term corpus, there were approximately 40 clusters identified by the function. For each graph, a standard score was calculated for each cluster. The score was used to determine clusters with majority female or male terms, where a larger positive value represented a cluster with majority terms with skew toward female representation and a larger negative value represented more terms with skew toward male representation:

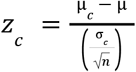

Where *μ*_*c*_ indicates the mean sex imbalance value for the cluster, *μ* indicates the mean of all sex imbalance values, *σ*_*c*_ indicates the standard deviation value for the cluster, and *n* represents the number of clusters.

We visualized these clusters by projecting the disease term embeddings for each disease term into a two-dimensional space using the t-distributed stochastic neighbor embedding (t-SNE) method. We originally generated term embeddings using BiomedBERT [37]. However, after projecting the embeddings onto the two dimensional space, similar disease terms did not colocalize as expected. To ensure similar diseases clustered appropriately, we introduced a hierarchical structure into our term corpus through mapping the original terms to the Mondo Disease Ontology [32] (see **Methods**, Mapping extracted disease terms to Mondo identifiers). We applied the Fisher transform and normalized the research-derived sex imbalance values before plotting to visually compare disease groups across plots, then overlaid the computed research-derived sex imbalance for each term onto the two-dimensional projections, resulting in visually distinct disease groups.

### Global sex-specific disease prevalence adjustment

Some diseases are sex-specific and therefore limited to a specific model (e.g., breast cancer, endometriosis, prostate cancer). To identify diseases with sex imbalance despite being prevalent in both males and females (e.g., cardiovascular disease, lung disease), and not limited to one model, we calculated “biological sex imbalance” by adjusting the research-derived imbalance values by the global male and female prevalence of each disease.

Both calculations of imbalance inform the overall disparity of female inclusion in biomedical research. Research-derived imbalance is useful for assessing imbalance directly derived from published literature, while biological sex imbalance is relevant to underrepresentation without known biological justification of female samples in scientific experiments.

To highlight individual disease terms with biological sex imbalance, we adjusted the research-derived values to incorporate the global prevalence of each disease in males and in females. The disease prevalence information was downloaded from the Global Health Data Exchange (GHDx) [38] resource from the Institute for Health Metrics and Evaluation (IHME) [39] on 12/8/2024. This resource provides male and female prevalence for 286 different diseases associated with 11 disease categories in the database (e.g., cardiovascular diseases, diabetes, neoplasms, etc). The disease terms extracted from metadata were mapped to the diseases using the same AI-supported workflow to map disease terms from study and publication metadata to Mondo identifiers (see **Methods**, Mapping extracted disease terms to Mondo identifiers). The associated research-derived imbalance values were adjusted to consider the prevalence. Specifically, imbalance values were adjusted as following: *Adjusted imbalance* = *research-derived imbalance* * (*min*(*F_prevalence* / *M_prevalence*)) / *max*((*F_prevalence* / *M_prevalence*)).

Where “F prevalence” is the female global prevalence of the disease, “M prevalence” is the global male prevalence, and “research-derived imbalance” is the sex imbalance value of the disease term (if multiple original terms were mapped to the same GHDx term, the sex imbalance values were averaged) before the prevalence adjustment. This method accounts for known biologically justified imbalance by reducing the magnitude of sex imbalance for terms with high prevalence in one sex. Disease terms with no match in the GHDx database were treated as unadjusted.

## Results

### Inferred sample sex labels are accurate and are not influenced by low quality samples

We visualized the ratio of female to male samples for each study (**Figure 3b, 3c**), calculated as: 2 * *F_samples* / (*F_samples* + *M_samples*) – 1. Negative inflection indicates majority male samples and a positive inflection indicates majority female samples. Our gold standard was purposefully created to have an equal representation of male and female samples (**Figure 3a**) [22]. Equal representation of male and female samples ensured that further analyses would not be impacted by unequal expression information for one sex in our gold-standard subset. Individual studies within the gold-standard sample set span a wide range of sex representation (**Figure 3b**).

**Figure 3.**
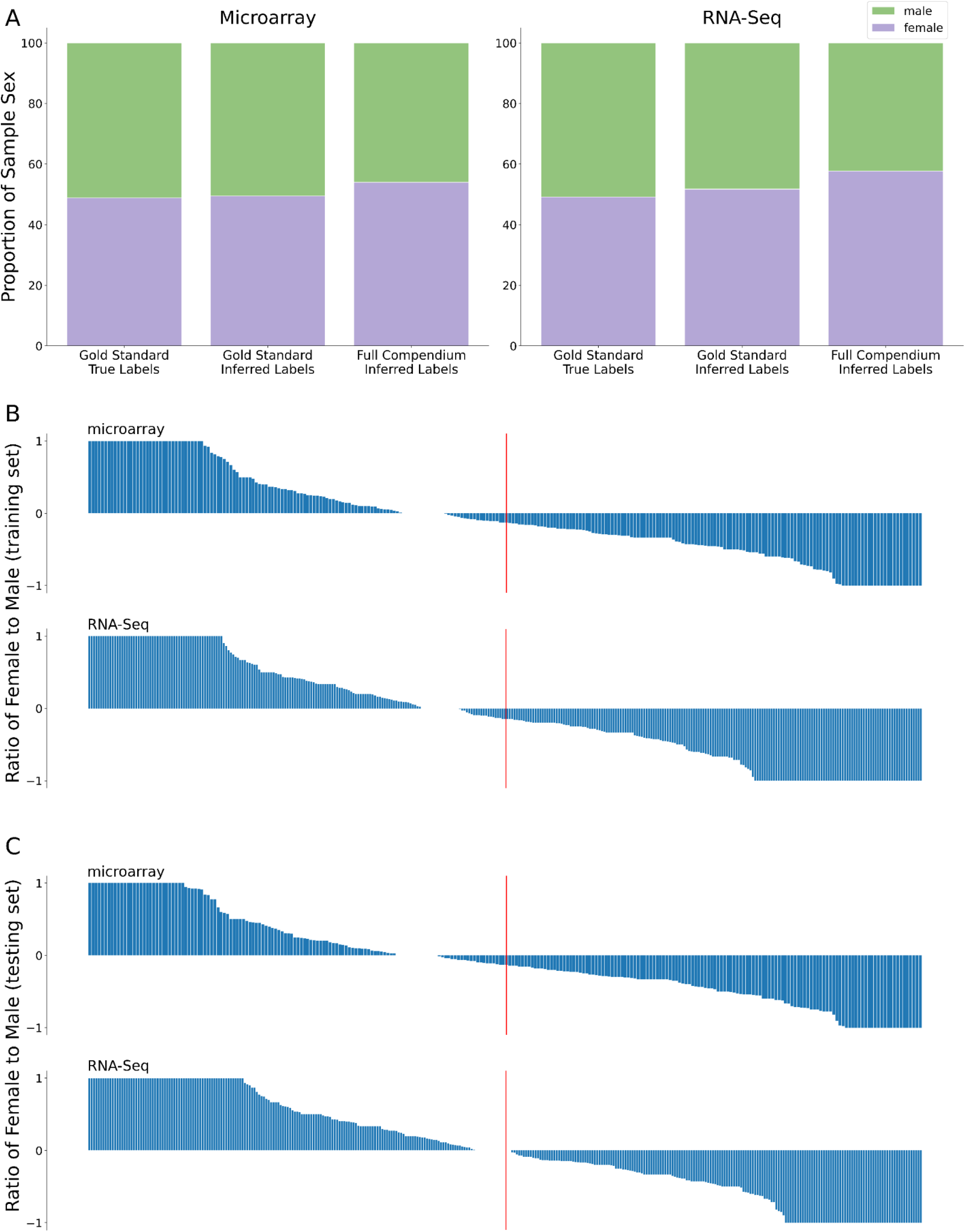
Proportion of male and female samples for manually labeled sex in gold standard subset, inferred sex in gold-standard subset, and inferred sex for entire sample compendium (A). Proportion of male and female samples in each study from GEO (microarray samples) and SRA (RNA-Seq samples) databases included in the manually curated gold-standard subset. Studies with a value of 1 have all female samples and studies with a value of -1 have all male samples (B). Study-level proportion of male and female samples in gold-standard subset after inferring sample sex (C). Red lines indicate 50% of studies.

To verify our sex label inference method, we inferred the sex of the ∼30,000 samples in our manually-curated subset. We used nine genes that demonstrated high accuracy at separating manually labeled samples based on expression values (separate processes for microarray and RNA-Seq samples) (**Figure 2b**). Combining microarray and RNA-Seq results, 97% of sample sex labels were correctly inferred (98% of microarray and 95% of RNA-Seq samples). Across both microarray and RNA-Seq samples, 4.5% of male samples were incorrectly inferred to be female and 1.8% of female samples were inferred to be male. Observing the study-level male and female representation, there was an increase in the percent of studies with majority female RNA-Seq samples (and decrease in the percent of studies with majority male RNA-Seq samples) after inferring the sex of each sample in the curated compendium (**Figure 3c**). Similarly, for studies with RNA-Seq samples, there were more studies with all female samples than all male samples after inference. In contrast, there were slightly more studies with majority male microarray samples after inferring labels and a similar percent of studies with all male and all female studies after inferring sex (**Figure 3c**). For both technologies, there was generally a similar study-level sex distribution compared to our expert curated sex labels (**Figure 3b**), further confirming the accuracy of our sex label inference method.

After verifying our method accuracy, we inferred the sex of our full transcriptome compendium of ∼230,000 human samples. There was a strong bimodal distribution of inferred sex for both microarray and RNA-Seq samples, with few samples near the decision boundary (**Figure 4**). Most samples were indicated as male or female by over 80% of the predicting genes, giving high confidence to our inferred sex labels.

**Figure 4.**
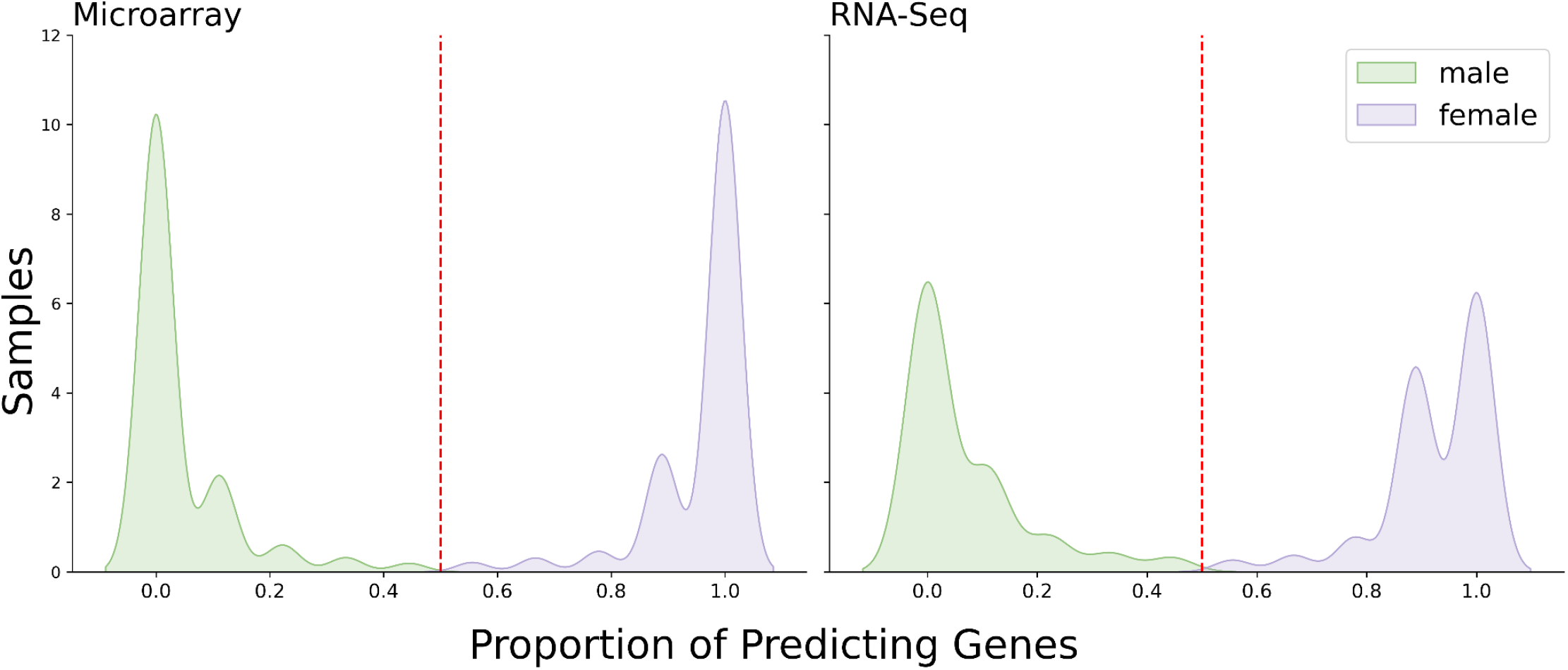
Distribution of predicted sample sex for microarray samples (left) and RNA-Seq (right) samples in the entire transcriptome compendium. Vertical red line indicates 50% of genes indicating a given sample as female (purple; above 50%) or male (green; below 50%).

The distribution of samples inferred to be female was higher than the number of samples inferred to be male, with 54% of microarray samples in the full compendium inferred as female compared to 49% in the gold-standard subset (**Figure 3a)**. For RNA-Seq samples, 57% of samples were inferred as female while 49% were labeled in the gold standard (**Figure 3a**). We were curious if this imbalance was due to increased representation of female subjects in our full sample compendium or a limitation of our inference method. We investigated whether poor sample quality led to the recorded expression levels to be lower than the actual values. The majority of genes used for inference are located on the Y chromosome (higher expression associated with male), so lower recorded expression level could lead to incorrect sample classification of a female label.

To investigate this hypothesis, we quantified the number of “poor quality” samples (see **Methods**, Inferring sample sex labels), and observed only few samples with low quality (i.e., high percent of genes with low expression compared to expression across all samples) (**Figure 5**). The majority of samples had few low expression genes, indicating that most samples are of good quality and poor sample quality does not explain the slightly increased representation of female samples in our inferred labels.

**Figure 5.**
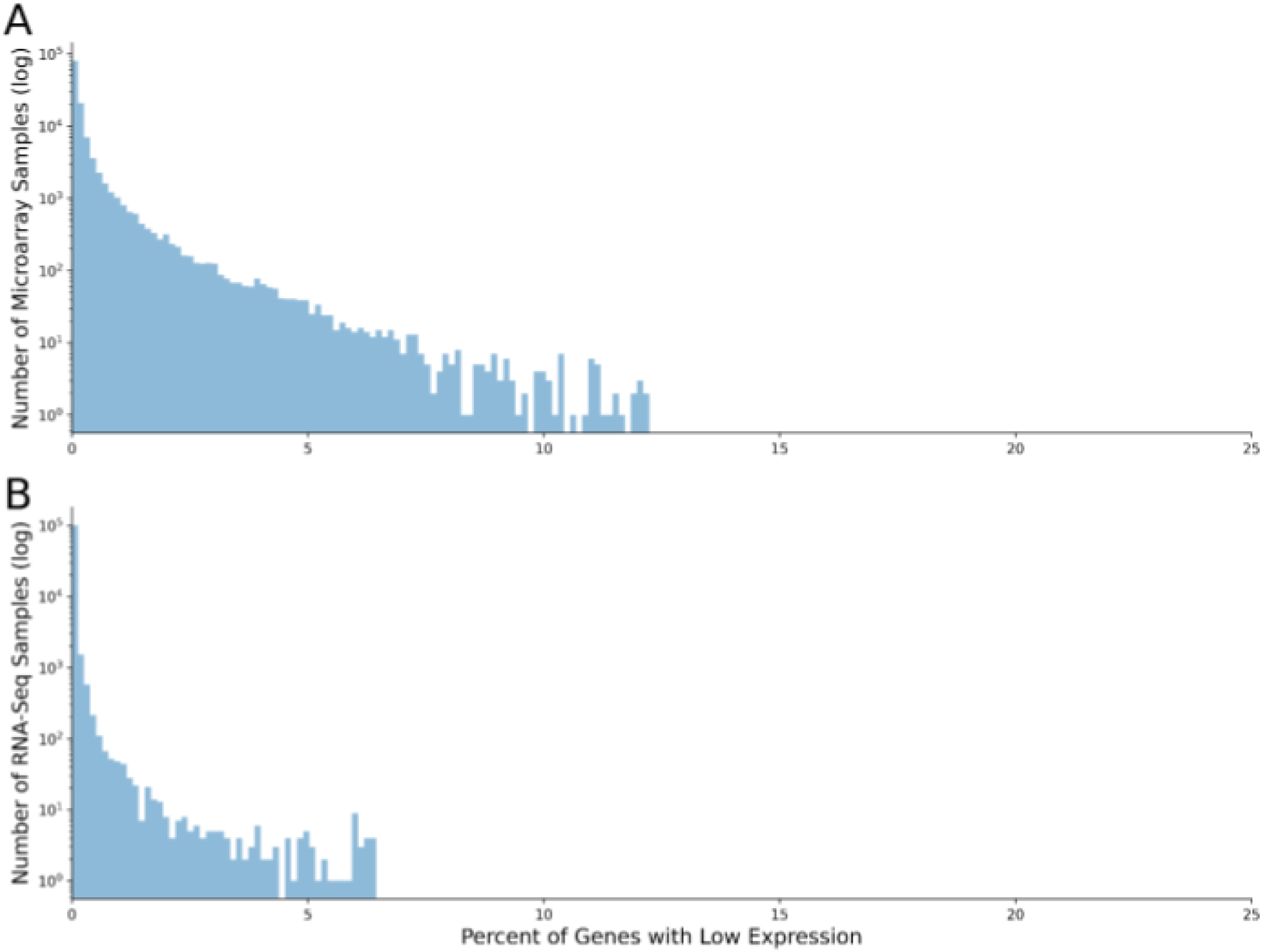
Distribution of microarray (A) and RNA-Seq (B) sample quality, defined as the percentage of genes with sex-balanced expression (i.e., genes expressed at similar levels in female and male samples) with low expression for each sample. Low expression was quantified as less than 25th percentile gene expression across all samples of the same predicted sex.

### Male skew is prevalent across disease terms with highest research-derived imbalance magnitude

To quantify sex imbalance across disease research areas, we extracted disease terms from the metadata of studies and publications associated with our ∼230,000-sample compendium, yielding between 1,825 and 6,176 terms depending on the metadata source and sequencing technology (see **Methods**, Disease term extraction from publication and study metadata). After mapping these terms to the Mondo Disease Ontology to enable cross-term comparisons, we calculated a research-derived sex imbalnce score for each term reflecting how strongly it co-occurred with male- or female-dominated studies. Across all four metadata corpora (microarray and RNA-Seq × studies and publications), imbalance scores spanned the full range from strong female to strong male skew, with terms skewed toward male outnumbering terms skewed toward female in every corpus even before adjusting for disease prevalence. Of the terms extracted from publication metadata, 61% (microarray-associated publications) and 54% (RNA-Seq associated publications) of the top 100 terms with the highest magnitude of imbalance were skewed toward male without known biological justification, while 17% (microarray) and 14% (RNA-Seq) of top disease terms skewed toward female with no known biological justification (**Supplemental Table 1**). From study descriptions, 44% (microarray) and 48% (RNA-Seq) of terms were associated with male overrepresentation and 23% (microarray) and 14% (RNA-Seq) of terms were associated with female overrepresentation. Examples of terms skewed toward male representation with manually-determined research-derived imbalance included “hepatocellular carcinoma,” “coronary artery disease,” “myocardial infarction,” and “inflammatory disease.” Examples of terms skewed toward female representation with research-derived imbalance included “cancer,” “colorectal cancer,” “pancreatic cancer,” and “metastasis.”

### Research-derived sex imbalance is identifiable for groups of disease terms

After manually investigating the sex imbalance of individual disease terms, we performed clustering analysis to observe distinct groups of disease terms with similar research-derived sex imbalance across our full disease term set with an automated approach. We calculated a standard score for each cluster (see **Methods**, Clustering of Mondo-mapped disease terms). A large positive score indicated that the cluster contained many terms associated with female overrepresentation. Examples of clusters with large positive scores included terms related to breast cancers, ovarian cancers, arthritis and joint diseases, bone diseases, multiple sclerosis, and thyroid cancers (**Figure 6**). A cluster with a large negative score indicated the cluster had more terms associated with male overrepresentation. Examples of clusters of male-skewed terms included terms related to prostate cancers, atherosclerosis and heart diseases, leukemia and lymphomas, mental health disorders, liver diseases, and lung diseases (**Figure 6**). The coherent grouping of disease areas by sex imbalance suggests that male and female underrepresentation in research are not scattered randomly across the disease taxonomy, but are concentrated in specific biological domains; a pattern not visible from individual term rankings alone.

**Figure 6.**
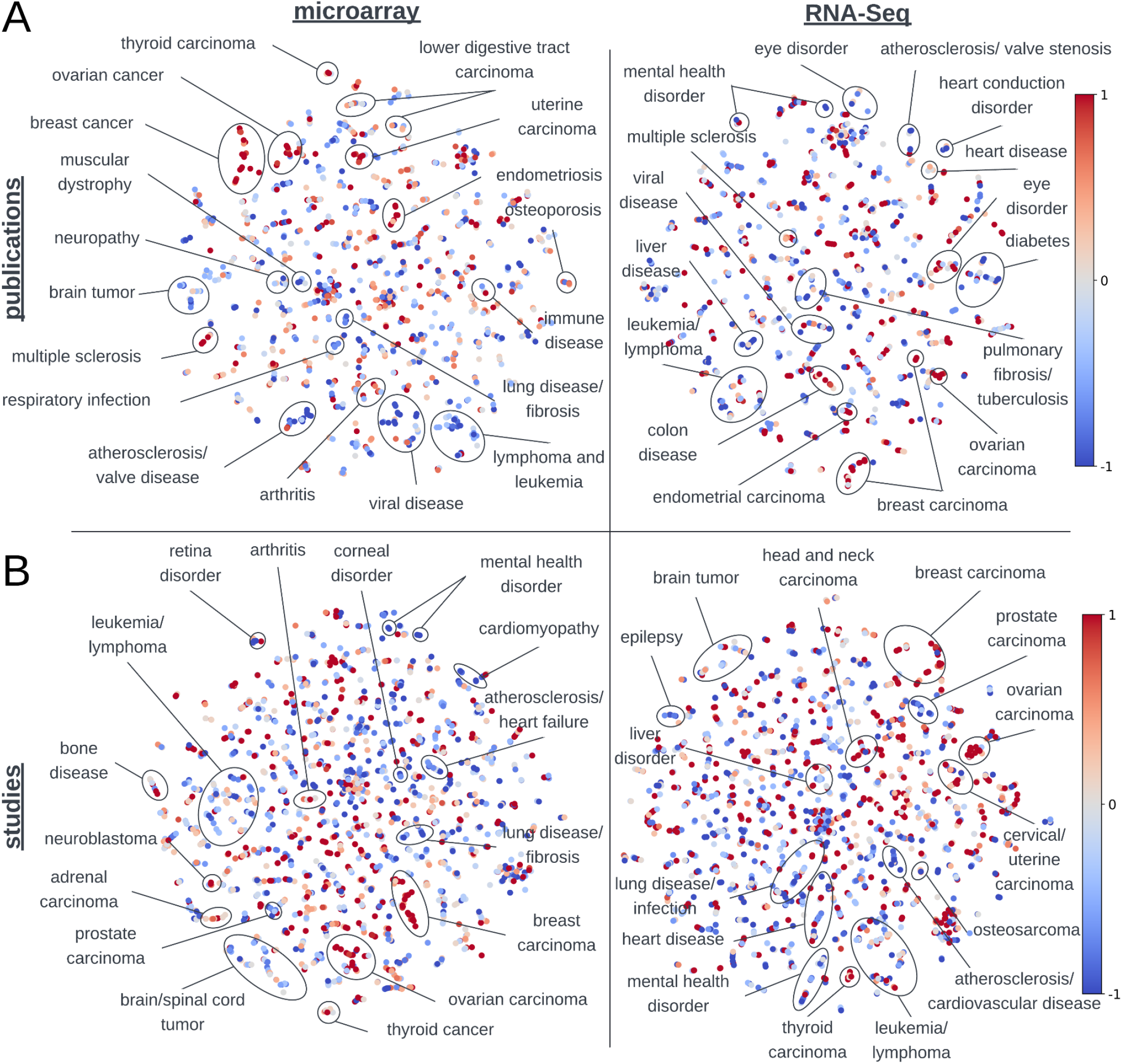
t-distributed stochastic neighbor embedding (t-SNE) of disease terms from publications (A) and studies (B) associated with microarray (left) and RNA-Seq (right) samples. Each point on the graph represents a Mondo disease term. Coloring indicates female (red; 1) or male (blue; -1) sex ratio correlation (research-derived sex imbalance).

### Adjusting for global, sex-specific disease prevalence to quantify biological sex imbalance

Calculating the sex ratio correlation for each disease term identifies the research-derived sex imbalance of the term across the full publication or study description corpus, but terms with the most severe imbalance do not reveal novel information as most have known biologically justified imbalance (e.g., “breast cancer” with female overrepresentation, “prostate cancer” with male overrepresentation). To draw out terms that do not fall into this category, we calculated the biological sex imbalance of each disease term by adjusting the research-derived imbalance value with the sex-specific global prevalence (**Supplemental Table 2**). The adjustment deflates the imbalance value magnitude proportionally to the degree of known sex-specific prevalence, so that diseases with equal prevalence in both sexes are unaffected while diseases with extreme prevalence skew are adjusted maximally toward zero. The new terms with the greatest magnitude of imbalance bring novel information about diseases that are understudied in a certain sex without known biological justification. For example, “breast cancer” originally had an imbalance value of 0.24, but with the prevalence adjustment, the new imbalance value was 0.00. “Psoriasis,” similarly prevalent in both males and females according to our resource, was adjusted to an imbalance value of -0.049 from -0.050 (**Figure 7**).

**Figure 7.**
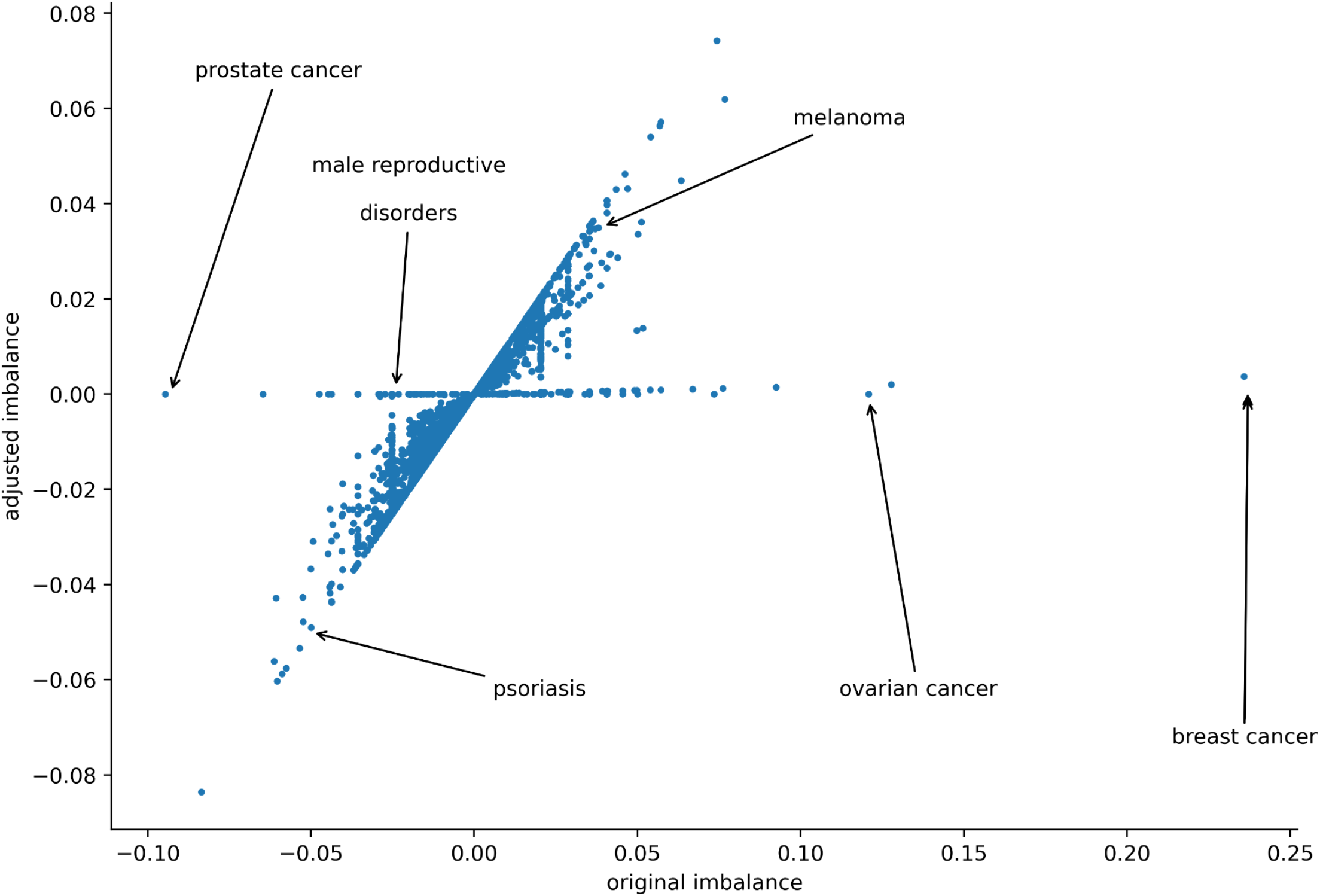
Sex imbalance adjustment of disease terms extracted from study descriptions associated with microarray samples. Adjustment accounted for global disease prevalence in males and females.

To visualize the effect of the prevalence adjustment and disease terms with the highest biological sex imbalance, we created word clouds displaying the top 100 terms with high imbalance values, where the size of each disease term is proportional to the term’s calculated sex imbalance value before and after adjustment (**Figure 8, Supplemental Figure 1**). Following the adjustment, many diseases with known sex-specific prevalence no longer dominated the highest-ranked terms, allowing diseases without obvious biological limitations to emerge. Assessing the top 100 terms, more terms were skewed toward male overrepresentation than female overrepresentation (**Figure 8, Supplemental Figure 1**). Terms with high biological male sex imbalance include “hepatocellular carcinoma,” “inflammatory disease,” “schizophrenia,” “myocardial infarction,” and “fibrosis” (**Supplemental Table 2**). Terms with high biological female sex imbalance include “uveal melanoma,” “pancreatic cancer,” “colorectal cancer,” and “intracranial aneurysms” (**Supplemental Table 2**). Adjusting the research-derived imbalance to incorporate the global prevalence of each disease identified terms that have higher sex imbalance without known biological justification.

**Figure 8.**
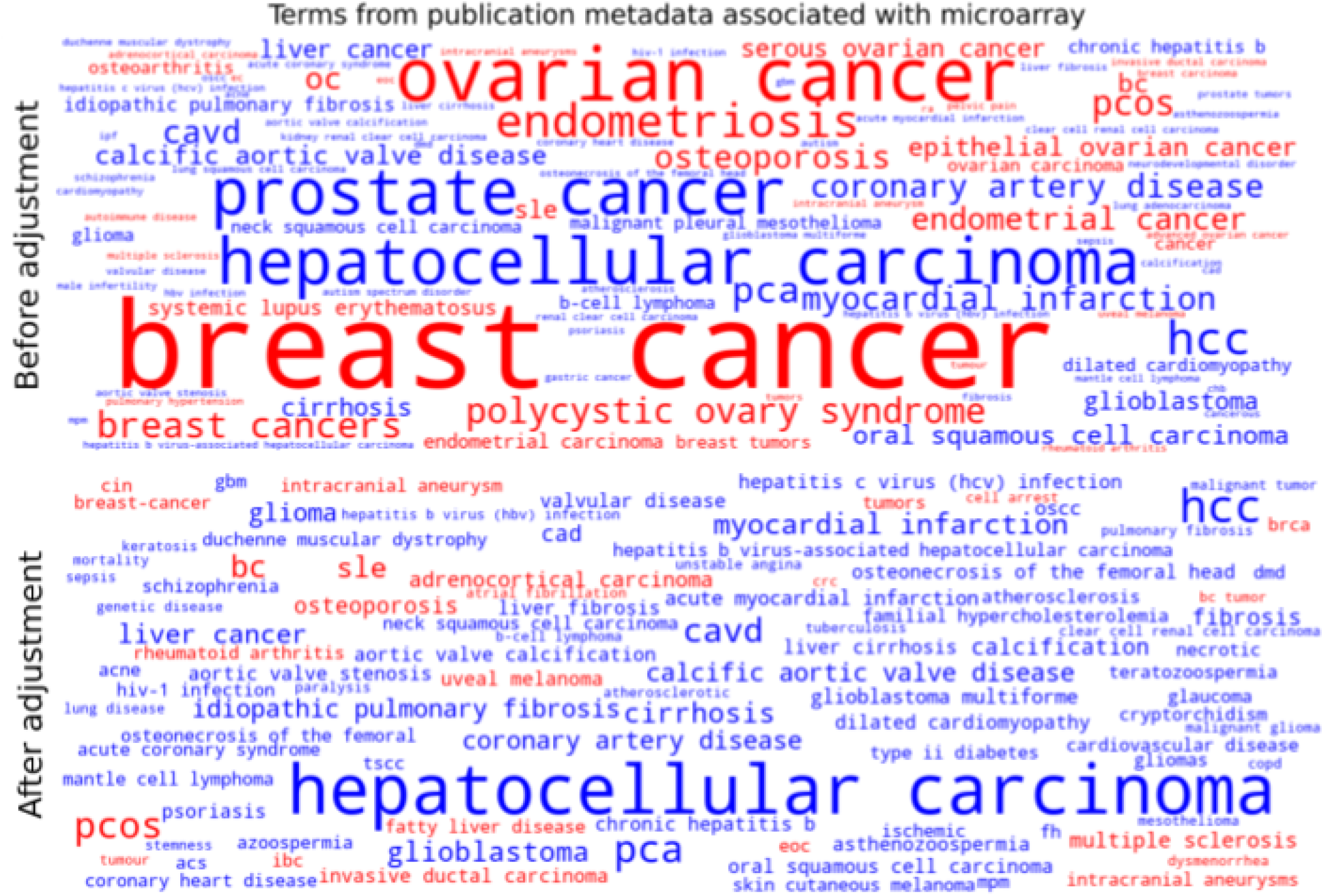
Word clouds of disease terms with highest absolute imbalance value (top 100 terms) before and after adjusting the sex imbalance. Terms from titles and abstracts of publications associated with microarray samples. Red word color indicates terms skewed toward female representation and blue word color indicates terms skewed toward male representation.

We further assessed biological sex imbalance across the full disease term corpus by calculating the percent of terms with adjusted sex imbalance above (female skew) and below (male skew) zero. We performed this calculation again, omitting terms with imbalance values close to zero (magnitude >0.001 and >0.01), in the case that the prevalence adjustment recalculated the sex imbalance value of terms with imbalanced prevalence to be near but not equal to zero. There was a higher percent of disease terms associated with male overrepresentation, where a higher or equal percent of terms were skewed toward male representation in almost every group (**Table 1**). For terms extracted from publication metadata, up to 58% of terms from publications associated with RNA-Seq samples and up to 57% of terms from microarray-associated publications were associated with male overrepresentation. For study metadata, up to 56% of terms from studies associated with microarray samples and up to 56% of terms associated with RNA-Seq samples were skewed toward male representation. This assessment indicates that even with adjusting imbalance values for global prevalence, a greater number of disease terms are associated with studies and publications that use a higher number of male subjects.

**Table 1.** Percent of terms with female (%f) or male (%m) biological sex imbalance. Terms extracted from either publication or study metadata associated with microarray or RNA-Seq biological samples. Percent of terms that skewed toward female or male representation was calculated for all terms, terms with an imbalance value greater than 0.01, and terms with an imbalance value greater than 0.001. A higher or equal percent of terms skewing toward male is indicated in bold.

|  | Microarray |  |  | RNA-Seq |  |  |
| --- | --- | --- | --- | --- | --- | --- |
|  | All terms | > 0.001 | > 0.01 | All terms | > 0.001 | > 0.01 |
|  | Publication |  |  |  |  |  |
| %f | 44% | 45% | 43% | 42% | 42% | 42% |
| %m | **56%** | **55%** | **57%** | **58%** | **58%** | **58%** |
|  | Study |  |  |  |  |  |
| %f | 47% | 47% | 47% | 44% | 44% | 50% |
| %m | **53%** | **53%** | **53%** | **56%** | **56%** | 50% |

## Discussion

Prior efforts that quantify male and female representation in biomedical research rely on studies that explicitly record participant sex or repositories where sex annotations are provided by individual researchers. This work establishes a different premise that sex imbalance is detectable at scale from the data itself by combining expression-based sex inference with ontology-grounded disease term extraction from study and publication metadata. As a consequence, the scope of auditable research expands from hundreds of manually examined publications to hundreds of thousands of samples linked to thousands of disease terms, shifting sex imbalance assessment from a manual monitoring task to a scalable, repeatable computational audit.

To make these estimations, we first inferred the biological sex of 229,528 human transcriptome samples based on each sample’s expression of 18,478 genes. Our method successfully inferred 98% of microarray and 95% of RNA-Seq samples, comparable to the 91-93% reported by Flynn et al. [19] using logistic regression on a larger but metadata-labeled sample set. The modest accuracy advantage likely reflects the higher sample quality enforced during our gold-standard curation [22,23]. Accuracy for the full ∼230,000 sample compendium cannot be directly assessed, as sex labels were not extracted for all samples. However, the low rate of poor-quality samples (**Figure 5**) supports the reliability of inferred labels across the compendium.

After inferring the sex for our full transcriptomics compendium, we estimated the research-derived (directly calculated from the correlation between terms and the ratio of female to male samples in study or publication metadata) and biological (accounting for global disease prevalence in each sex) sex imbalance of disease terms extracted from the metadata of 8,704 studies and 5,021 publications associated with the samples. Our approach identifies disease terms related to male-(“prostate cancer”) or female-specific organs (“breast cancer,” “ovarian cancer”), confirming the capability of our method to find groups of terms with their expected sex association.

Further, our method confirms previous findings of diseases with male or female research-derived sex imbalance, indicating one sex to be understudied for that disease without accounting for sex-specific disease prevalence. Male imbalance has been reported in studies related to nervous system drugs [19], male organ-related disease terms (“prostatic neoplasms,” “testicular germ cell tumor”), and cardiovascular diseases [17]. We report male research-derived sex imbalance in similar areas, including cardiovascular and heart disease, vascular disorders, brain tumors/neurodevelopment disorders, and epilepsy, as evidenced by high correlation with imbalanced sample distribution toward male subjects. For females, we report high imbalance values in female organ-related disease terms (ovarian and uterine carcinomas, endometriosis), autoimmune diseases, and bone-related disorders, also in agreement with previous reports [17,19].

After incorporating male and female global disease prevalence, we report related terms such as “viral infection,” “schizophrenia,” “myocardial infarction,” “hiv infection,” “liver cancer,” and “glioblastoma” to have high male biological sex imbalance. Sex imbalances in these disease research areas could have serious ramifications. For example, the protective effect of estrogen against glioblastoma progression [40] positions post-menopausal women at higher risk than their age-matched male counterparts. In schizophrenia, females develop more severe side effects than males, despite demonstrating better treatment compliance [41]. Sex plays a critical role in disease treatment response. In combination with understudied female cohorts and, as a result, treatment development optimized in male-dominated datasets, female patients are potentially at higher risk for negative outcomes.

Lastly, our approach discovers new disease areas with sex imbalance in the current biomedical research space based on samples included in studies and publications. Areas such as eye disorders, leukemia and lymphomas, lung diseases and pulmonary fibrosis, and liver diseases were identified to have research-derived male imbalance. Specific terms related to these areas were identified to have high male-skewed biological imbalance after incorporating global sex-specific disease prevalence, including “leukemia,” “acute lung injury,” “lung diseases,” “pulmonary fibrosis,” “liver cancer,” and “liver disease.” We also identified terms with female-skewed biological sex imbalance, including digestive system cancers (“pancreatic cancer,” “colorectal cancer”), bone diseases (“osteoporosis,” “sarcoma”), and immune system diseases (“multiple sclerosis”). However, across all collections of terms, more disease areas were quantified to be associated with male overrepresentation without known biological justification (**Figure 6, Figure 8**).

We acknowledge the following limitations of our study. The increase in the proportion of female samples in our inferred labels is inherent to our sex label inference method. We hypothesized that this phenomenon was due to low sample quality, which can result in altered recorded expression levels [42]. Eight of nine genes used in the sex label inference method are located on the Y chromosome (**Figure 2b**). Higher expression was associated with male samples for most of our predicting genes, so low sample quality could result in an inferred female label for a sample. Our results suggested this possibility, as more labeled male samples were inferred as female (2.5% of microarray samples, 6.9% of RNA-Seq samples) than labeled female samples inferred as male (1.3% of microarray, 2.4% of RNA-Seq) in our gold standard inferred labels. However, we investigated the quality of our samples by comparing the gene expression for each sample to the expression for all samples, and found few samples to be of poor quality (**Figure 5**). Two alternative explanations remain unresolved. Cell line samples (which are abundant in public repositories) can lose Y-chromosome through generations of subculturing [43], causing male-derived cell lines to be misclassified as female. Alternatively, the full compendium may indeed contain more female-derived samples than the deliberately balanced gold-standard subset. Whether this skew introduces directional bias in the downstream sex ratio estimates (and, if so, in which direction) cannot be determined without sex labels for the full compendium. Distinguishing these possibilities would require cross-validation against independently derived metadata annotations such as those in MetaHQ [23].

Disease terms extracted from study and publication metadata included terms that are relevant to diseases but not diseases themselves, such as “adult,” “biosynthesis,” “damage,” and “invasive.” Furthermore, some extracted terms included acronyms such as “cavd,” “pcos,” “pca,” and “soc.” Our model extracts biomedically-relevant disease terms and ensuring all extracted terms are verified diseases would be a time-consuming manual effort, or limit the flexibility of the model to extract a broad range of specific terms. As a result, some terms such as these might not be mapped properly to terms in the Mondo Disease Ontology and those in the GHDx database, which may affect their reported sex imbalance values. For these mappings, we used a custom, AI-supported workflow. The limited capability of the AI model, whose failure modes (conservative matching, occasional hallucination) might cause some terms to be miscategorized or excluded, may effect downstream clustering and prevalence-adjusted results. Additionally, some matchings were slightly different from the target term (e.g., incorrect capitalizations, spelling, truncated phrases), which had to be manually corrected. However, with the advancement in AI technology, it is likely future models will have improved context to correctly handle abbreviations, disease-related terms, as well as produce fewer mistakes in the output. An alternative productive next step would be further developing the RAG pipeline and AI API call to manage these cases.

Lastly, our approach combining TF-IDF values and inferred sex labels through Spearman correlation assumes a monotonic relationship between the relevance of a disease term to a given document and the sex ratio of the term’s associated samples (i.e., terms appearing in male-dominated studies should consistently appear more often in male-dominated studies, not just sometimes). If that monotonicity breaks down for certain disease areas (e.g., terms that appear in both studies with a high proportion of male subjects and those with a high proportion of female subjects for different biological reasons), the correlation might underestimate the true sex imbalance. The result of this possibility might limit our approach in reporting all disease areas with significant imbalance, however, it would further promote the areas we do report as research areas potentially at risk for a detrimental lack in sex-specific knowledge.

In conclusion, we assessed male and female sex imbalance in the current biomedical research space on a large-scale, inferring the sex of ∼230,000 human transcriptome samples and extracting relevant biomedical disease terms from ∼5,000 studies and ∼9,000 publications associated with the samples. Before this work, auditing sex imbalance in the biomedical literature required either manual inspection of a small number of journals or publications, or relied on self-reported demographics (i.e., sex of study subjects) that are absent for most publicly available samples. The framework presented here enables such an audit computationally across hundreds of thousands of samples and thousands of disease terms, and to repeat it as the public data landscape grows. Researchers, funders, and journals can use the ranked disease term outputs to identify where compensatory studies are most urgently needed and to track whether investment in those areas is closing the gap over time.

## Declarations

### Ethics approval and consent to participate

All data used in this study is publicly available from national databases (https://www.ncbi.nlm.nih.gov/geo/, https://www.ncbi.nlm.nih.gov/sra).

### Consent for publication

Not applicable.

### Availability of data and materials

The datasets analyzed in the current study are available at https://github.com/krishnanlab/dis-sex-imbalance included in the supplemental files, or available from the corresponding author upon reasonable request.

### Competing interests

The authors declare that they have no competing interests.

### Funding

This work was primarily supported by US NIH NIGMS grant R35 GM128765 and NSF grant 2328140 to A.K.

Authors’ contributions

L.E.V. and A.K. designed the study. L.E.V. and H.Y. developed the software. L.E.V., K.A.J., and M.A. curated data. L.E.V. performed the analyses. L.E.V., P.H., H.Y., M.A., and A.K. interpreted the results. L.E.V. wrote the final manuscript with feedback from P.H., H.Y., M.A., and A.K. All authors read and approved of the final manuscript.

## Acknowledgements

We would like to thank Keenan Manpearl for providing materials used in analyses, and all members of the Krishnan Lab for valuable feedback on the study.

## Notes

https://github.com/krishnanlab/dis-sex-imbalance

### Competing Interest Statement

The authors have declared no competing interest.

