## Supplemental Figure 1 for "Large-scale automated detection reveals pervasive sex imbalance in biomedical research"

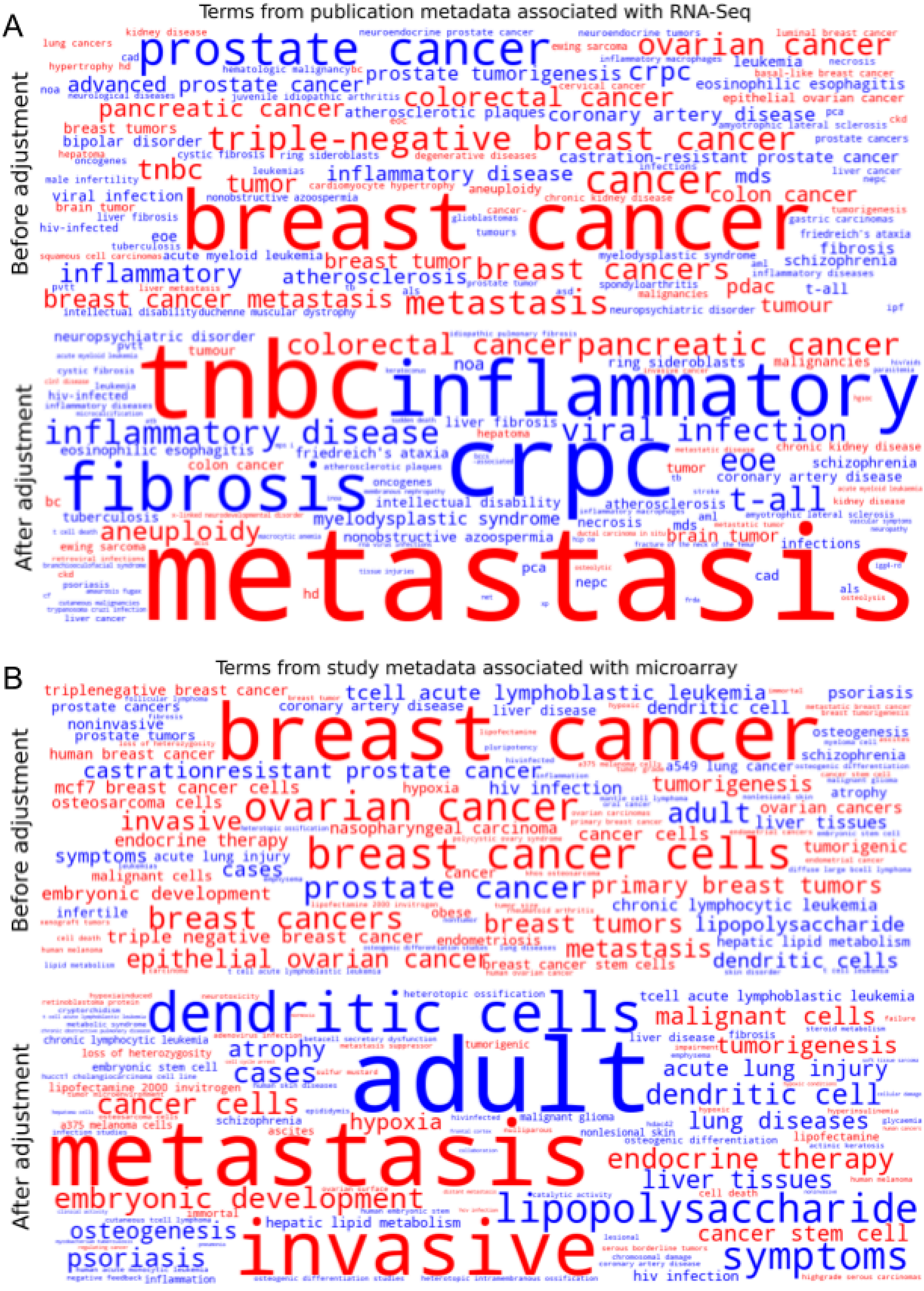
A
Terms from publication metadata associated with RNA-Seq
Before adjustment
prostate cancer
ovarian cancer
breast cancer
triple-negative breast cancer
colorectal cancer
pancreatic cancer
advanced prostate cancer
prostate tumorigenesis
crpc
coronary artery disease
castration-resistant prostate cancer
tnbc
cancer
tumor
inflammatory disease
mds
colon cancer
viral infection
inflammatory
atherosclerosis
breast tumor
breast cancers
breast cancer metastasis
metastasis
pdac
tumour
leukemia
eosinophilic esophagitis
epithelial ovarian cancer
bipolar disorder
breast tumors
schizophrenia
fibrosis
t-all
acute myeloid leukemia
myelodysplastic syndrome
friedreich's ataxia
After adjustment
tnbc
inflammatory
crpc
fibrosis
metastasis
colorectal cancer
pancreatic cancer
inflammatory disease
viral infection
eoe
t-all
aneuploidy
myelodysplastic syndrome
nonobstructive azoospermia
brain tumor
eosinophilic esophagitis
friedreich's ataxia
neuropsychiatric disorder
ring sideroblasts
liver fibrosis
schizophrenia
coronary artery disease
atherosclerosis
intellectual disability
tuberculosis
oncogenes
infections
malignancies
B
Terms from study metadata associated with microarray
Before adjustment
breast cancer
tcell acute lymphoblastic leukemia
castrationresistant prostate cancer
ovarian cancer
breast cancer cells
prostate cancer
primary breast tumors
breast cancers
breast tumors
lipopolysaccharide
epithelial ovarian cancer
metastasis
dendritic cells
dendritic cell
adult
invasive
tumorigenesis
tumorigenic
mcf7 breast cancer cells
osteosarcoma cells
hypoxia
hiv infection
endocrine therapy
nasopharyngeal carcinoma
symptoms
acute lung injury
cases
embryonic development
chronic lymphocytic leukemia
triple negative breast cancer
hepatic lipid metabolism
triplenegative breast cancer
prostate cancers
human breast cancer
psoriasis
osteogenesis
schizophrenia
a549 lung cancer
atrophy
ovarian cancers
liver tissues
cancer cells
malignant cells
endometriosis
After adjustment
dendritic cells
adult
metastasis
invasive
lipopolysaccharide
symptoms
malignant cells
tumorigenesis
acute lung injury
dendritic cell
lung diseases
endocrine therapy
liver tissues
embryonic development
cancer stem cell
osteogenesis
psoriasis
cancer cells
atrophy
cases
hypoxia
tcell acute lymphoblastic leukemia
chronic lymphocytic leukemia
loss of heterozygosity
embryonic stem cell
lipofectamine 2000 invitrogen
hepatic lipid metabolism
hiv infection

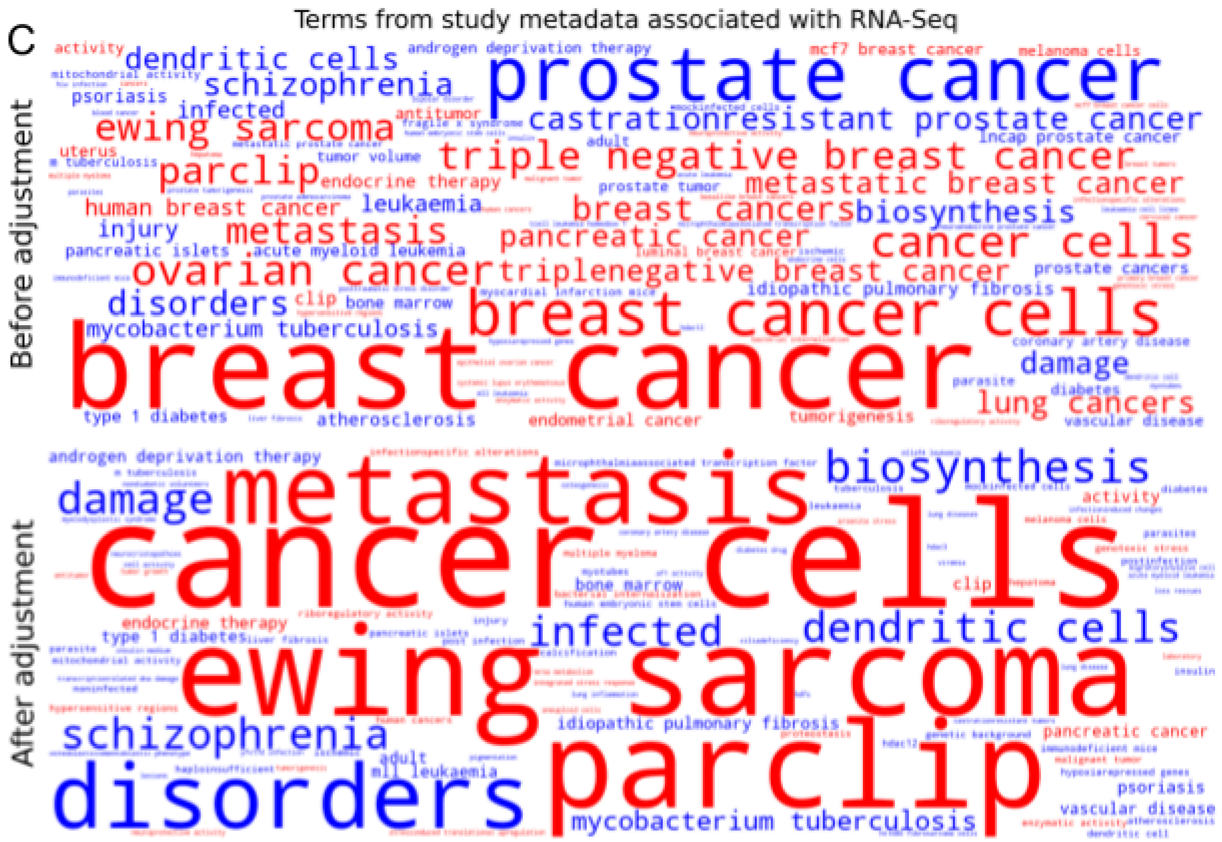


**Supplemental Figure 1.** Terms from titles and abstracts of publications associated with RNA-Seq (A) samples and from descriptions of studies associated with microarray (B) and RNA-Seq (C) samples. Red word color indicates terms skewed toward female representation and blue word color indicates terms skewed toward male representation.
